# Mechanical counterbalance of kinesin and dynein motors in microtubular network regulates cell mechanics, 3D architecture, and mechanosensing

**DOI:** 10.1101/2021.05.25.445700

**Authors:** Alexander S. Zhovmer, Alexis Manning, Chynna Smith, James. B. Hayes, Dylan T. Burnette, Alexander X. Cartagena-Rivera, Rakesh K. Singh, Erdem D. Tabdanov

**Author notes:** **Correspondence authors**: Alexander S. Zhovmer, Rakesh K. Singh, (**Lead contact**): Erdem D. Tabdanov. These authors contributed equally (Alexis Manning, Chynna Smith).

## Abstract

Microtubules (MTs) and MT motor proteins form active 3D networks made of unstretchable cables with rod-like bending mechanics that provide cells with a dynamically changing structural scaffold. In this study, we report an antagonistic mechanical balance within the dynein-kinesin microtubular motor system. Dynein activity drives microtubular network inward compaction, while isolated activity of kinesins bundles and expands MTs into giant circular bands that deform the cell cortex into discoids. Furthermore, we show that dyneins recruit MTs to sites of cell adhesion increasing topographic contact guidance of cells, while kinesins antagonize it *via* retraction of MTs from sites of cell adhesion. Actin-to-microtubules translocation of septin-9 enhances kinesins-MTs interactions, outbalances activity of kinesins over dyneins and induces discoid architecture of cells. These orthogonal mechanisms of MT network reorganization highlight the existence of an intricate mechanical balance between motor activities of kinesins and dyneins that controls cell 3D architecture, mechanics, and cell-microenvironment interactions.

## INTRODUCTION

Microtubules (MTs) and MT motors have emerged as key drivers of dynamic changes to the cell cytoskeleton. They control cell chirality ^1,2^, shape plasticity ^3–7^, 3D architecture ^8–11^, motility signaling ^12^, mechanotransduction ^13^, and 3D migration (in immune cells) ^14^. In cooperation with the integrin-based actomyosin adhesion system, MTs serve as a mechanosensor that integrates extra- and intracellular mechanical signals on a scale of the whole cell ^15^. Mechanistically, MTs form a cellular network of unstretchable rod-like cables that are embedded within a sterically confining, laterally re-enforcing actomyosin networks providing cells with means to resist mechanical deformations ^16,17^ - *e*.*g*., in cardiac myocytes, MTs that are coaxially integrated into the myofibrillar network act as rod-like springs which mechanically oppose myofibril contractile shortening ^18,19^. In other cell types, MTs provide scaffolds that mechanically reinforce actively protruding pseudopodia, *e*.*g*., invadopodia ^7,20–22^, podosomes ^23,24^, invasive microtentacles ^25,26^, or dendritic cell protrusions that facilitate cell adhesion and spreading within soft fibrous 3D collagen matrices ^27,28^.

Cell 3D architecture is a result of a dynamic balance between the internal and external structural, biochemical, and mechanical signals that are integrated by actomyosin and MT networks. For instance, cell body linearization is driven by an actomyosin-dependent inward collapse of the MT network into linear MT bundles ^10,29^, whereas integrin-dependent cell adhesion and spreading drives MT unbundling ^30–32^. Adaptations of cell shape to migration, *e*.*g*., dynamic interactions between the cell and the extracellular matrix (ECM), also require integration of biochemical and biomechanical signals between MTs and actomyosin which enables cells to sense, align with, and exhibit guided motility along nanotextured 3D microenvironments ^32^. While it is clear that MT and actomyosin networks play a crucial role in the control of cell shape and cell-microenvironment interactions, the precise nature of specific interactions between MTs and actomyosin, and the manner in which mechanical forces are transmitted and balanced between these two networks, remains elusive.

An accumulating body of evidence indicates that MT motor proteins actively control cell architecture and mechanics *via* regulating interactions between MTs and other cellular components like actomyosin and organelles as well as with the extracellular matrix (ECM). For example, recent findings demonstrate that dynein motors drive and coordinate spatial alignment of radially organized MT networks, centering MTOCs at the geometrical or mechanical center of the actomyosin network of the cell ^33,34^. Dyneins also control cell-type-specific changes in the cell architecture that drive downstream functional adaptations; *i*.*e*., in neurons, dyneins drive MT sliding and extension along the axonal axis ^35–38^, while in hematopoietic cell lineages, dyneins generate tensile forces in MT networks that surround the cell nucleus and ultimately result in the formation of nucleus-compressing 3D MT meshworks that favor mechanosensory activation of chromatin towards myeloid (and away from lymphoid) differentiation ^39^. Meanwhile, kinesins demonstrate similar mechanical and MT-restructuring effects, but in apparent opposition to the actions of dyneins. For example, kinesins have been shown to promote bundling and sliding of antiparallel, “telescopic” MT filaments that manifests as “looping” at the level of individual filaments ^40^ and drives global reorganization of MTs at the level of the whole cell ^41^. These individual findings highlight the involvement of both kinesin and dynein motors in the control of cell structure and shape. However, very few studies, if any, have probed how or whether the synergy or antagonistic actions of both motors in tandem drive changes in cell shape, mechanics, or cell-microenvironment interactions, presumably due to challenges associated with cytotoxicity and the general lack of motor-specific inhibitors to study mesoscopic (whole cell) scale effects of kinesin and dynein balance.

Previous studies have linked dynamics of actomyosin and MT networks to large-scale changes in cell structure ^9,42^ using MT networks of enucleated mammalian platelets that have MTs organized into parallel, 2-4 µm ring-like structures known as marginal bands ^43^. During platelet activation, the action of dynein drives band expansion into the actomyosin platelet cortex ^9,44^, resulting in displacement of tension into the cortex, platelet discoid deformation, and eventually a disc-to-sphere transition ^8,45^. Thus, the 3D structure of platelets is dependent upon a delicate, mechanistic balance between antagonistic dynein-based MT forces and actomyosin-based tension within the platelet cortex that can be dynamically shifted *via* modulating dynein activity or that of other motors like myosins ^8,9,45^. As the direct mechanical antagonist of dyneins, kinesins oppose and prevent dynein-driven marginal bands overextension ^44^, likely *via* crosslinking/bundling MTs in the opposite direction of dynein-dependent MT antiparallel sliding ^9,42^. However, the exact mechanism of how marginal band homeostasis is maintained by the opposing actions of dynein-kinesin remains unresolved; *i*.*e*., whether the dyneins and kinesins drive marginal band rings extension-compression balance *via* the indirect MT-actomyosin-MT ^9^ or the direct MT-MT filaments crosslinking and sliding ^42^. Moreover, the exact roles and mechanisms for dyneins and kinesins during platelets’ marginal band structural condensation around a singular multi-coiled MT ^43^, remain unknown.

Despite the significant progress in understanding of isolated effects for MT motor proteins on the organization of MT network and MT-actomyosin interaction, the importance and extent of mechanical antagonism between dyneins and kinesins at the mesoscopic whole cell level, *e*.*g*., for a cell with a regular size and unreduced structural complexity, remain elusive. It is also unclear which role the balance between kinesin and dynein motors activities plays in the cellular mechanobiology and cell-microenvironment interactions, *e*.*g*., structural adaptation and migration responsiveness to the microenvironment cues.

## RESULTS AND DISCUSSION

### Inhibition of dyneins and myosins induces giant microtubular rings

First, we examine the morphological response of MDA-MB-231 triple negative human breast adenocarcinoma cells following inhibition of dyneins and myosins. For efficient dyneins inhibition we utilize Dynapyrazole A, a Ciliobrevin-derived inhibitor, which displays substantially higher stability and dynein-inhibition efficacy than that of *per se* Ciliobrevins ^46^. Similarly, we utilize (-)-Blebbistatin for suppression of actomyosin contractility ^9,32^. To capture the in-tissue complexity of cell adhesion faithfully, yet uniformly, we use an isotropic rectangular grid of collagen-1 “fibers” that support fibrillar cell spreading^11^ **(Figure 1a)**. Collagen type-1, a major cell adhesion ligand in the ECM of physiological tissues, is organized into orthogonal sets of 1 µm-wide parallel lanes, spaced with 15 µm-wide pitch to mimic collagen “fibers”. Micropatterned collagen grids prevent the excessive formation of the 2D lamellipodium ^11,47^. Abnormal 2D lamellipodium dominates cell adhesion and spreading scenarios on regular flat substrates, while 1D “fiber” architecture induces a more natural quasi-3D modality of cell-ECM interactions^47^. In order to closely recapitulate the average tissue mechanical rigidity, we micro-print the collagen ECM grids on polyacrylamide hydrogel surfaces with a mechanical rigidity of G’=50 kPa ^48,49^.

**Figure 1.**
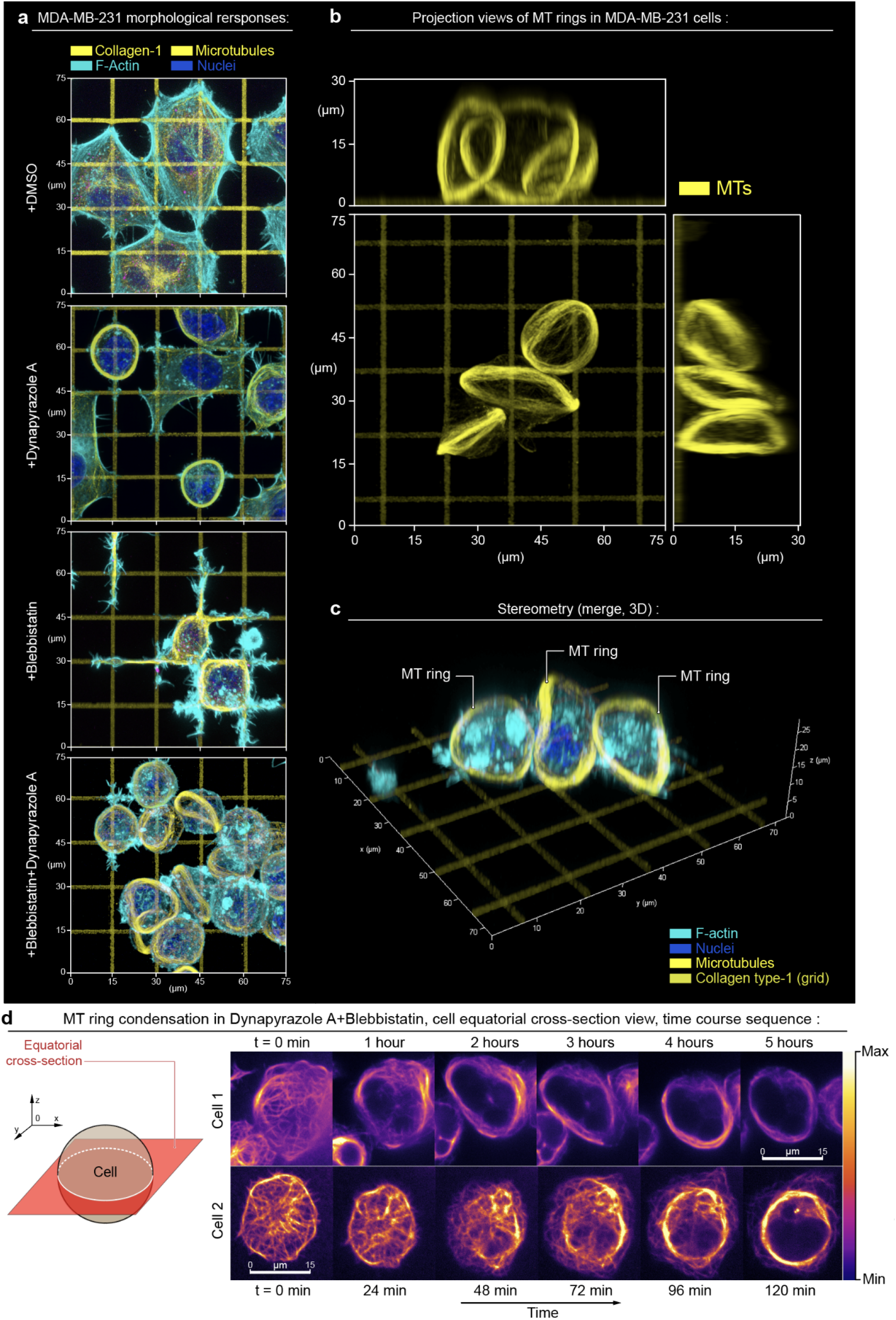
Top 3D overview of MDA-MB-231 cells during adhesion and spreading along the artificial extracellular matrix (orthogonal type-1 collagen grid, G’=50 kPa). **(a)** - Micrographs present cell configurations in the control condition **(+DMSO)**, during dynein inhibition **(+Dynapyrazole A)**, in the presence of actomyosin contractility inhibitor **(+Blebbistatin)**, and under combined dynein and actomyosin contractility suppression **(+Blebbistatin+Dynapyrazole A)**. Upon biaxial spreading along the orthogonal collagen lanes in the control condition **(+DMSO)**, cells develop concave polygonal architecture with stress-fibers predominantly located at the mechanically tensed cell periphery - *i*.*e*., arching free cell edges. Suppression of the dynein motor activity **(+Dynapyrazole A)** induces mixed cell architectures, with 70% of cells partially developing MT rings along with cells having polygonal architecture. Inhibition of actomyosin contractility **(+Blebbistatin)** shifts cell organization towards the spreading network of dendritic protrusions that compliantly extend along the collagen grid. These protrusions feature the microtubule scaffold as their structural core. Notably, combined dynein and myosin II inhibition **(+Blebbistatin+Dynapyrazole A)** results in the reorganization of the MT network into MT ring in >90% cells. Similarly to marginal bands in mammalian platelets, these rings deform cells into lenticular discoids. Separate channels are shown in **SI1**. **(b)** - 2D projection views of the MT rings. **(c)** - Stereometric 3D view, merged channels. **(d)** - Time course sequence for MT network expansion and ring condensation in MDA-MB-231 cells **(+Dynapyrazole A+Blebbistatin)**, equatorial confocal plane imaging, SiR-actin live F-actin labeling, see **Movie 1 and 2**. Note the MT network expansion-driven translocation from throughout the cell volume towards the cell cortex.

Within two hours, in control conditions **(+DMSO)**, spreading cells acquire concave polygonal shapes with stress-fibers accumulating at the arching cell edges **(Figure 1a, SI1, +DMSO)**. In this scenario, we observe a geometric configuration of adhesions, which presumably serves to induce high linear tension along free cell edges ^11,50^. Suppression of actomyosin contractility with Blebbistatin results in polygonal cell shape collapse and ‘dendritic’-like cell spreading with cell protrusions growing along the collagen-1 grids **(Figure 1a, SI1, +Blebbistatin)**, reflecting Arp2/3-driven low-contractility or ‘fluid-like’ cell spreading, reported for 2D/3D ECM ^11,27,28^. Dynein inhibition with Dynapyrazole A induces global reorganization of the MT network into MT bundles for ∼70% of cells, and circularized MT bundles (rings) in ∼50% of all cells **(Figure 1a, SI1, Dynapyrazole A)**. Combined inhibition of dyneins and myosins drives substantially more prominent and uniform bundling of MTs across the entire cell population, resulting in the loss of cell spreading and MT reorganization into 3D circular ring-like bundles (diameter Ø≈20 µm) that are located at the cell peripheral circumference **(Figure 1a, SI1, +Blebbistatin+Dynapyrazole A)**. 3D projections (X0Y, X0Z, Y0Z) for Blebbistatin+Dynapyrazole A-treatment **(Figure 1b,c, SI2)** indicate that MT rings induce deformation of cells into lenticular discoids (Ø≈20-25 µm). These discoids are structurally reminiscent of significantly smaller (Ø≈3 µm) MT marginal bands (MB) in highly specialized, enucleated, structural complexity-reduced mammalian platelets ^42^, and in non-mammalian red blood cells ^4^. However, these structures were not previously reported for mesenchymal cells, *e*.*g*., for large cells with unreduced structural complexity. Dynamics of MT ring formation in Dynapyrazole A+Blebbistatin shows that MTs, which are normally distributed roughly equally throughout the cell volume, translocate and re-distribute almost entirely into the cortex, indicating the active spatial expansion of MT network **(Figure 1d, Movies 1 and 2)**. Spatial expansion of MTs results in their spherical buckling into the MT ring, caused by the confining boundaries of the cell cortex **(Figure 1c)**. This phenomenon, largely similar, but not fully identical to the MB formation in much smaller mammalian platelets ^8,45^, highlights the mechanical antagonism between the expanding MT ring and the cell cortex.

### Balance of dynein and kinesin activities controls structure of microtubule network

MDA-MB-231 cells develop giant MT rings in response to the suppression of dynein activity, while the formation of MT marginal bands in mammalian platelets, on the contrary, requires active dyneins, yet inactive kinesins ^9^. Therefore, we hypothesize that the reorganization of the isotropic MT network into circular bundles is likely driven by the disbalance of dynein and kinesin motor activities in either direction **(Figure 2a-1)**. In this model, the action of dyneins and kinesins together creates an expansion-compaction balance of the MT network within the cell volume, as well as MT protrusion-retraction balance within cell adhesion sites **(Figure 2a-1)**. Isolated activity of kinesins (*e*.*g*., **+Dynapyrazole A**) leads to both MT network expansion and kinesin-driven retraction of MTs from cell adhesion sites **(Figure 2a-2)**. Physical expansion of MT network and retraction of MTs from peripheral adhesions results from kinesin-driven antiparallel sliding of MTs towards their distal (+)-ends, that, if uncompensated for by opposing dynein-dependent MT sliding, leads to decreased MT overlap and elongation of antiparallel MT bundles.

**Figure 2.**
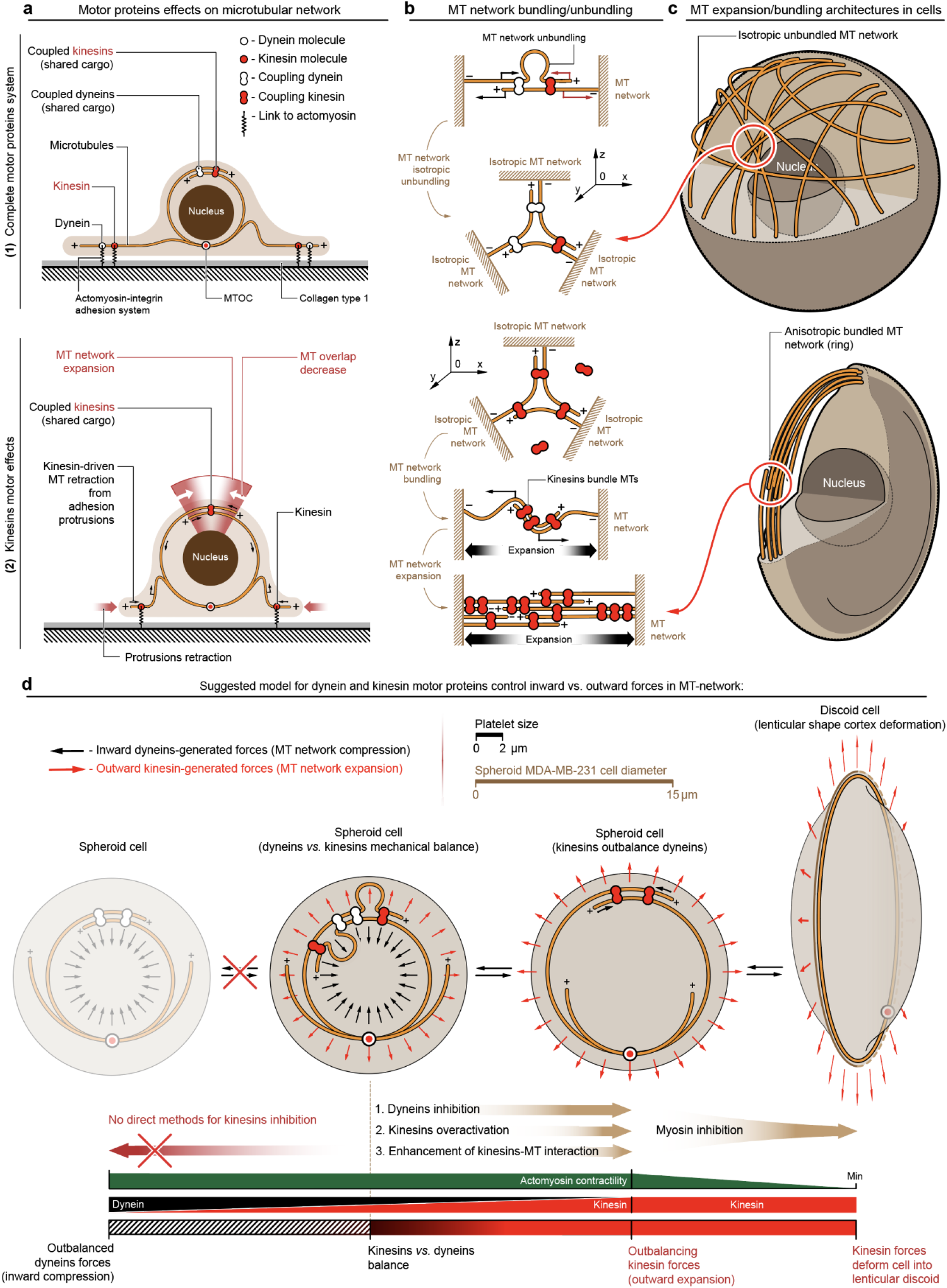
Kinesin and dynein motor proteins counterbalance outward and inward-directed forces within MT network. **(a)** - Schematic representation of dynein-kinesin superposition and sole kinesin-generated forces within the MT network: (**1**) - Superpositioned dynein and kinesin forces counterbalance each other within both cell volume and at cell-substrate adhesion sites. (**2**) - Kinesin motor proteins facilitate MT network expansion into the cell cortex within the cell volume *via* antiparallel telescopic MT sliding-out. Kinesin-driven MT sliding decreases MT overlap, extends overlapping antiparallel MT filaments (MT network expansion), and drives MT network retraction/expulsion from the cell-ECM adhesion sites. **(b)** - Key force-generating configurations of MT-motor proteins can effectively drive MT network compaction/expansion and MT bundling/unbundling. Note that dynein and kinesin motors both directly and indirectly bundle MTs, and form coupled complexes, *e*.*g*., by cargo shared between two or more motors (coupled dynein and kinesin). (**1**) - Simultaneous work of coupled dyneins and kinesins within antiparallel MT structures induce directionality conflict within sliding MTs, that results in MT filaments buckling and unbundling into scattered, misaligned and isotropic MT networks. (**2**) - Kinesin-only system (coupled kinesins) induces MT bundling and antiparallel MT ‘telescopic’ sliding-out that drives 3D MT network collapse into the ring (circular bundle) and ring expansion into the cell cortex deforming cells into the discoids. **(c)** - Non-treated spheroid cells with active kinesins and dyneins have unbundled MT network, scattered throughout cell volume. Isotropic MT network highlights dynamic mechanical balance of network compaction and expansion. Dynapyrazole A+Blebbistatin-treated MDA-MB-231 cells develop kinesin-bundled MT rings that feature kinesin-driven ring expansion. MT ring expansion deforms individual cells into lenticular discoids. **(d)** - Suggested mechanism describes mechanical and structural contribution of MT network into the cell cortex as a counterbalancing system of dynein and kinesin motors. Three alternative strategies are available for shifting the mechanical balance towards a higher contribution of kinesins - direct dynein inhibition **(+Dynapyrazole A)**, kinesin overactivation **(+Kinesore)**, and MT network redecoration with the septin-9 that enhances kinesin-MT interactions **(+UR214-9)**. Note that inhibition of cortex contractility, combined with outbalanced activity of kinesins, leads to the MT ring formation-expansion, and cell deformation into the lenticular discoid.

Notably, dyneins ^51^ and kinesins ^52–54^,55 each can crosslink and linearize MT networks into antiparallel bundles ^40^. These bundles are additionally enhanced through interaction with the actomyosin cytoskeleton which serves in the mechanical transmission of dynein-driven forces between adjacent MT filaments ^56^. Our basic idea of mechanical antagonism between dyneins and kinesins is that these motors exhibit individual and combined structuring effects onto the bundling and unbundling of MT networks **(Figure 2b)**. In the antiparallel MT bundles, the combined activity of dyneins and kinesins will lead to a MT sliding-in and sliding-out directionality conflict, consequent buckling of MTs which is followed by MT isotropic disarray ^57,58^, MT network unbundling and spatial misalignment **(Figure 2b-1)**. Alternatively, in parallel (unidirectionally-oriented) MT bundles, the principal lack of the antiparallel MT sliding activity will lead to low levels of MT reorganization and bundling dynamics for both dynein and kinesin motors. Thus, we neglect the contribution of the MT motor activity within the parallel MT bundles. Importantly, shifting the balance of dynein and kinesin motor activity *via* dominance of either motor enhances MT bundling due to directionally coherent sliding of MTs, and decrease of the MTs sliding directionality conflict, as demonstrated on the example of kinesin-wise balance shift **(Figure 2b-2)**, driving reorganization of the isotropic MT network into the single bundled structure **(Figure 1a, SI1: +Dynapyrazole A and +Blebbistatin+Dynapyrazole A, Figure 1b and SI2)**. Since there is no technically reliable method for a detectable kinesin inhibition in non-platelet cells, the MTs reorganization effects, caused by the shift of the dynein-kinesin balance towards the dynein activity in MDA-MB-231 cells are depicted only hypothetically **(Figure 2d)**.

Based on our model, we explain the formation of giant MT rings **(a)** and cell discoids **(b)** in Dynapyrazole A by the inhibition of dyneins and significant outbalance towards the kinesin-driven effects. These effects, such as MT bundling, extension of MT bundles and their expansion into the cell cortex, result in the formation of the giant MT rings. These rings generate outward-directed forces sufficient for discoid deformation of the low-contractility actomyosin cell cortex during experiments with +Dynapyrazole A+Blebbistatin co-treatment **(Figure 1a-c: +Dynapyrazole A+Blebbistatin; Figure 2d)**. Experimental characterization of the kinesin-driven expansion of MT networks into the MT rings during **+Dynapyrazole A+Blebbistatin** co-treatment demonstrates discoid deformation and identifies robust accumulation of total kinesins within the MT rings **(Figure 3a, SI3)**, along with substantial MT ring diameter Ø dominance over the cells transverse width **(Figure 3b, SI3a,b)**. Notably, our data indicate that suggested mechanism of MT network reorganization by the balance of dynein and kinesin motor activities is more universal than could be previously anticipated based on the mammalian platelets cytoplast model ^9^, as the MT ring formation and the cell discoid deformation take place within various cells featuring significantly larger scale and complexity **(Figure 2d)**.

**Figure 3.**
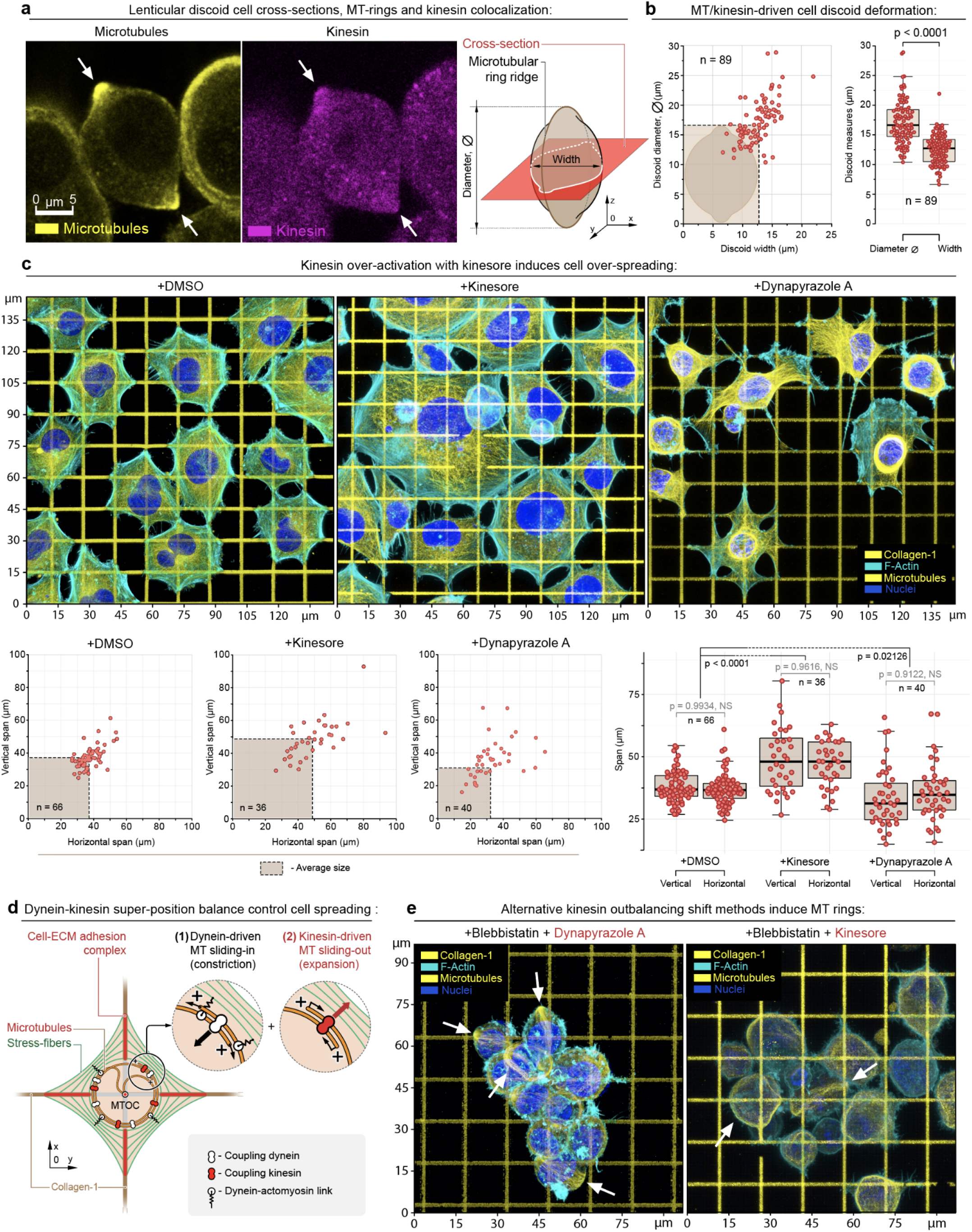
Structural analysis of the MDA-MB-231 cells during alternative dynein-kinesin balance shift mechanisms. **(a)** - MT ring median-plane cross-section in MDA-MB-231 cells (*left*) as shown in the schematic (*right*) in **+Dynapyrazole A+Blebbistatin**. Note kinesin accumulation at the MT rings (*arrows*), see also **SI3a-d**. **(b)** - MT rings expand and deform cells into the lenticular discoids (*MT ring diameter > cell width*). **(c)** - *Top row* - Adhesion and spreading of MDA-MB-231 cells on the collagen-1 grids (G’=50 kPa) in control condition **(+DMSO)** and during kinesin overactivation **(+Kinesore)**, compared to the direct dynein inactivation **(+Dynapyrazole A)**. Kinesin over-activation with **+Kinesore** does not inhibit the dynein activity, yet induces a significant increase of the biaxial spreading of cells in comparison to the control cells. Alternatively, dynein-kinesin balance shift towards the kinesin activity *via* the direct dynein inhibition **(+Dynapyrazole A)** induces a visible reduction of cell spreading and “dendritization” of cells that is similar the loss of cell contractility shape during Blebbistatin-induced myosin II inhibition **(Figure 1, +Blebistatin)**. The loss of cell contractility in Dynapyrazole A is related to the two main factors ^59^: the loss of MT-to-actomyosin mechanical crosslinking and force transmission, as well as the loss of the dynein-driven mechanical contractility within MT network **(+Dynapyrazole A)**. *Bottom row* - Morphometric analysis of control **(+DMSO)**, and **+Kinesore**- and **+Dynapyrazole A**-treated cells indicate a significant increase in cell biaxial spreading, driven by kinesin-mediated MT network expansion. **(d)** - Suggested mechanism for the MT-actomyosin interaction in cells on the collagen grids. Dyneins and kinesins control the **(1)** MT network compaction and **(2)** MT network expansion respectively. Dyneins also crosslink the MT network to the actomyosin cytoskeleton (stress-fibers). *I*.*e*., actomyosin, adhesion-bound matrix, adhesion complex, MTs and MT motor proteins subsystems are united into a single mechanical system. See also **SI4a**. **(e)** - Alternative methods for outbalancing the kinesin activity *via* **+Dynapyrazole A+Blebbistatin** and **+Kinesor+Blebbistatin** induce MT rings and cell discoids. Pairwise one-way *t* tests-derived p values are shown on the plots with corresponding n (size of individual cells measurement sets) for each condition, generated in triplicates.

### Control of cell-ECM adhesion and spreading by dynein-kinesin balance

We next proceed to examine the effect of the superposition of dynein and kinesin motor activities, that form a dynamic and mechanically antagonistic balance, on cell adhesion and spreading on collagen ECMs. To do so, we modulate the dynein-kinesin balance *via* two alternative strategies: (a) kinesin overactivation with Kinesore^40^ and (b) kinesin-wise shift *via* Dynapyrazole A-induced dynein suppression. Remarkably, Kinesore-driven kinesin overactivation enhances polygonal cell spreading along orthogonal collagen-1 lines compared to the controls **(Figure 3c, +DMSO** *vs*. **+Kinesore)**. Alternatively, kinesin-wise balance shift *via* Dynapyrazole A-induced dynein inhibition leads to dendritization and poor polygonal spreading **(Figure 1a and Figure 3c, +DMSO** *vs*. **+Dynapyrazole A)**. Dendritization is analogous to the low cell contractility spreading mode seen during Blebbistatin treatment **(Figure 1a, SI1, +Blebbistatin)**. We attribute these differences to the dynein activity status - *i*.*e*., MT network contraction-expansion balance is controlled by the dynein-kinesin mechanical balance **(Figure 3d, SI4a)**. However, dyneins facilitate mechanical cross-linking and mechanotransduction of the forces within the MT network (**SI5a**) and the actomyosin cytoskeleton ^33,35,57,59–62^ **(Figure 3d)**. Thus, suppression of dyneins with Dynapyrazole A reduces both the MT network contractility (**SI5c**) and the MT-actomyosin mechanotransduction leading to the partial loss of the overall cell-ECM contractility and impaired cell spreading. Therefore, we propose that a mechanical synergy between actomyosin and MT-dynein contractile machineries in the adhesion sites **(Figure 3d, SI4a)** facilitates the mechanosensory-stimulated cell adhesion and spreading ^11^ **(SI4a-1)**. In this case, Kinesore-induced cell over-spreading **(Figure 3c, +Kinesore)** is a result of the Kinesore-enhanced kinesin activity that expands the MT network and preserves background contractility of the actomyosin-dynein-MT system, which sustains cell adhesion and spreading *via* mechanosensory stimulation.

Notably, the non-spreading cell behavior, induced by the combination of kinesin over-activity and low actomyosin contractility, results in the formation of MT rings and discoid cells regardless of dynein activity status **(Figure 3e, +Kinesore+Blebbistatin, +Dynapyrazole A+Blebbistatin, SI3e-f)**. Our results indicate that MT rings are a structurally ‘self-locked’ system that requires only MT-to-MT filaments bundling and mechanical sliding, *e*.*g*., by kinesins ^52–55^ **(Figure 2a-b, 3)**. That is, it does not require a dynein-mediated mechanical or structural integration with actomyosin for MT rings formation, as it does during cell adhesion-spreading on ECM. Furthermore, the dispensable role of dynein activity during the kinesin-driven MT ring formation, both in +Kinesore+Blebbistatin and +Dynapyrazole A+Blebbistatin conditions, indicates that MT rings are the extreme of kinesin outbalance within the existing range of dynein-kinesin mechanical balances, as both treatments induce the dynein-kinesin balance shift towards the kinesin activity *via* two principally different mechanisms.

### Microtubule translocation of septin-9 induces MT ring formation

Recent studies report that translocation of septin-9 onto the MTs enhances kinesins activity, *e*.*g*., kinesin-3/KIF1A ^63^. We adopt this mechanism as an alternative strategy for kinesin-wise dynein-kinesin balance shift **(Figure 4a)**. First, we confirm the septin-9 translocation from actomyosin stress-fibers onto the MT network upon septin-2 inhibition with UR214-9 **(Figure 4b, SI6a)**, a high-e ciency non-cytotoxic septin-2 inhibitor ^64,65^, recently derived from cytotoxic forchlorfenuron (FCF) septin inhibitor ^66^. Inhibition of the septin-2, a binding core of septin filaments, induces disintegration of the septin complexes ^67^, and UR214-9-induced septin-9 release and translocation onto the MT network, followed by the kinesin overactivation **(SI6b)** formation of MT rings in the MDA-MB-231 cells **(Figure 4b, +UR214-9)** similarly to **+Dynapyrazole A±Blebbistatin (Figure 1a)**, and to **+Kinesore+Blebbistatin (Figure 3e)**. Notably, UR214-9 does not show the signs of dyneins suppression **(SI6c)**, but induces MT network bundling-expansion into the MT rings *via* the septin-9-kinesin pathway **(SI4b)**. Thus, this orthogonal strategy confirms that the outbalance of kinesins over dyneins is a su cient requirement for the MT network bundling and expansion into emergent ring-like structures, regardless of the particular molecular pathway leading to kinesins mechanical outbalance (**SI5b** and **SI6b**). The advantage of this approach is that UR214-9 allows stimulation of kinesins on MTs regardless of the background cellular activity of kinesins and dyneins (**SI4b** and **SI6b,c**).

**Figure 4.**
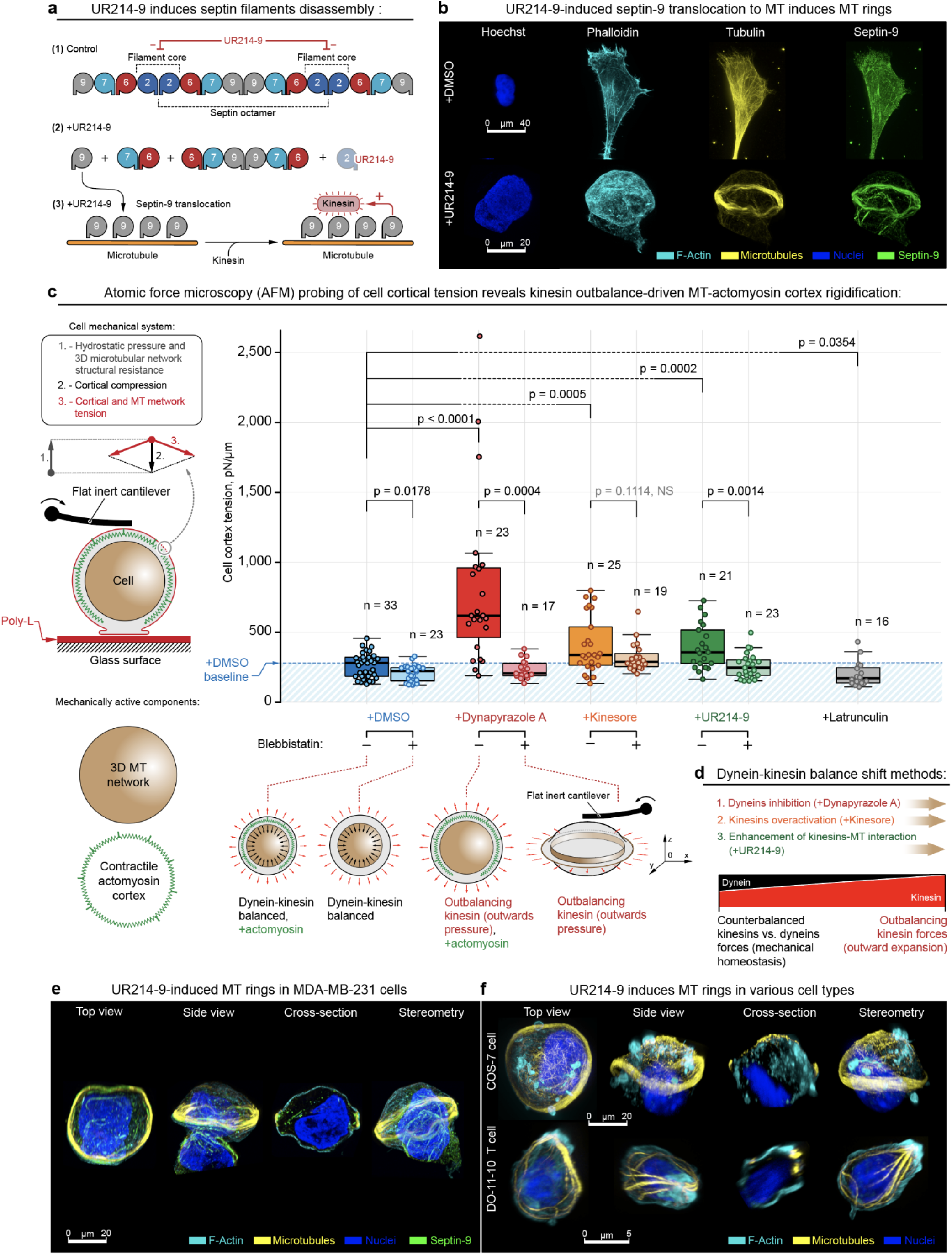
Formation of MT ring *via* alternative dynein-kinesin balance shift mechanisms features cell rigidification as an outcome of MT network expansion. **(a)** - UR214-9-targeted septin-2 inactivation leads to actin-to-microtubules translocation of septin-9, followed by enhancement of kinesin-MT interaction, and increased kinesin activity. **(b)** - Immunofluorescent visualization of stress-fiber-specific localization for septin-9 in MDA-MB-231 cells in control conditions **(+DMSO)**, and septin-9 translocation onto MT network, accompanied by MT ring formation during septin-2 inactivation **(+UR214-9)**. See also **SI6a**. **(c)** - AFM analysis of the MDA-MB-231 cells rigidification upon kinesin-wise activity outbalance, induced by three alternative mechanisms. *Left* - Schematic of the cell mechanical system: both MT network and actomyosin represent mechanically active cell components that synergistically regulate cell mechanics. *Right* - Dynapyrazole A, Kinesore, and UR214-9 induce cell rigidification. *Bottom* - schematic representation of cell rigidification upon kinesin’s outbalancing activity: while **+Dynapyrazole A+Blebbistatin, +Kinesin+Blebbistatin**, and **+UR214-9+Blebbistatin** all induce MT rings, due to the rings’ anisotropic configuration their mechanical effects are non-readable by AFM. **(d)** - Schematic representation of the alternative mechanisms for inducing the kinesin’s outbalancing activity. Both dynein inhibition **(+Dynapyrazole A)**, kinesin over-activation **(+Kinesore)**, and kinesin-MT interactions enhancement *via* septin-9 F-actin-to-MT translocation **(+UR214-9)** unanimously induce cell rigidification *via* kinesin-driven MT network expansion. See also **SI4b**. **(e, f)** - UR214-9-induced kinesin overactivation results in MT rings formation in MDA-MB-231 cells (**e**), as well as in other tested cell lines, *e*.*g*. - COS-7 (**f**, *top*), and in DO-11-10 murine T cell line (**f**, *bottom*). Pairwise one-way *t* tests-derived p values are shown on the plots with corresponding n (size of individual cells measurement sets) for each condition, generated in triplicates.

### Cell mechanics is controlled by dynein-kinesin balance

We summarize the described above mechanistic findings into a cell mechanical model that predicts kinesin-driven cell rigidification upon kinesin-wise dynein-kinesin balance shift, followed by the MT network expansion **(Figure 1d, Movies 1 and 2)**. *I*.*e*., the MT network mechanically expands into the cell cortex, providing the cortex with the additional, outward-directed force-generating MT scaffold **(Figure 2a-3, SI4b)**. This model-based prediction is strongly supported by cell discoid deformation with the expanding MT ring into the relaxed actomyosin cell cortex in Blebbistatin **(Figure 3a-b, SI3a-b)**. We test the predicted kinesin-wise outbalance-induced cell rigidification with atomic force microscopy (AFM) live cell probing **(Figure 4c)**. Indeed, we identify a significant cell rigidification (described by increase in surface cortical tension) across all three methods of the kinesin-wise MT motors balance shift. *I*.*e*., both **+Dynapyrazole A, +Kinesore**, and **+UR214-9** treatments unanimously induce the MDA-MB-231 cells rigidification **(Figure 4c, SI4b)**. Their combinations with Blebbistatin reduce AFM rigidity readout to a lower level, due to actomyosin cortex relaxation **(a)** and MT network collapse into the spatially anisotropic ring **(b)**, which is largely unavailable for AFM cantilever probing **(Figure 4c)**. Thus, indeed all alternative and orthogonal methods of kinesin-wise MT motors balance shift induce MT scaffold structural expansion, mechanistically and structurally proving the existing dynein-kinesin mechanical balance that orchestrates MT network architecture and cell mechanics **(Figure 4d, SI4b)**.

The universal character of changes in cell structural organization and its resulting mechanics, induced by three alternative mechanisms of dynein-kinesin activities balance shift **(Figure 4c, SI4b)** also extends into the universality of described MT networks structural reorganization in various cell types. *I*.*e*., UR214-9-induced MT bundling-expansion into MT rings are observed not only in the principal MDA-MB-231 cell model **(Figure 4e)**, but appear to be universal for otherwise dissimilar cell types, such as COS-7 (∼20% of cell population, 2 hours of UR214-9 incubation period) and DO-11-10 murine T cells (∼15% of cell population, 2 hours of UR214-9 incubation period) **(Figure 4f)**, highlighting the basic and universal importance of septins-MT-motor proteins signaling axis. Thus, the MT-motor proteins rebalancing effects are fundamental to cell mechanobiology regulation principles.

### Dynein-kinesin balance controls 2D contact- and “2.5D” steric-cell guidance

Dynein motors regulate structural coherence between MT and actomyosin networks ^33,36,68^. On the other hand, MT network mechanically (sterically) scaffolds cellular contact guidance on nanotextures ^32^. Therefore, we hypothesize that tunable coupling of MTs and actomyosin within the cell adhesion sites by balancing dynein-kinesin motor activity may play a significant role in cellular contact guidance at the adhesion sites **(Figure 2a)**. According to this model, the sole kinesin-driven MT network ‘self-locking’ into the MT ring and MT retraction from actomyosin cell adhesion interface can be in direct competition with the dyneins that facilitate mechano-structural integration of MT filaments into the cell actomyosin adhesion system ^33^, and away from the giant MT rings structures **(Figure 2a)**. Notably, this model is complementary to the previously reported indispensable scaffolding role of MT networks for cell adhesion and spreading ^20^ by supporting low-contractility dendritic cell protrusions ^11,27,28^.

We propose that dynein activity facilitates the integration of MT filaments as the contact guidance scaffold into the cell adhesion and into spreading actomyosin protrusions **(Figure 5a, +DMSO)**. This activity of dyneins competes with antagonistic kinesin activity **(Figure 5b-1)**, rendering dyneins as the key MT motor subsystem that controls cellular contact guidance *via* mechanical coupling of MTs and actomyosin, and by pulling MT filaments into the nano-grooves as the cell spreading-elongation-directing scaffold. Indeed, Blebbistatin-induced suppression of actomyosin contractility does not affect cell contact guidance elongation along the nanotexture, as the MTs in-groove alignment remains intact **(Figure 5a, c, +DMSO** *vs*. **+Blebbistatin)**. On the contrary, Dynapyrazole A-induced dynein inhibition completely abrogates insertion and elongation of MTs into and along nano-grooves on collagen-1 nanotextures, which is accompanied by the complete loss of cell alignment to the contact guidance cues of nanotextures **(Figure 5a and 5c, +DMSO** *vs*. **+Dynapyrazole A)**. Additionally, kinesin activity outbalance induced by UR214-9 leads to identical results, such as the loss of contact guidance and formation of planar MT rings that are unaligned with underlying nanotextures **(Figure 5a and 5c, +DMSO** *vs*. **+UR214-9)**. Instead, MT network reorganizes into the ‘self-locked’ bundled configurations with uncompensated kinesin activity that also retracts and expels the MTs from the actomyosin cytoskeleton in cell adhesion sites **(Figure 5b-2)**. In summary, dynein-kinesin balance controls cellular contact guidance in a sterically interactive microenvironment by recruitment of MTs to the sites of cell adhesion.

**Figure 5.**
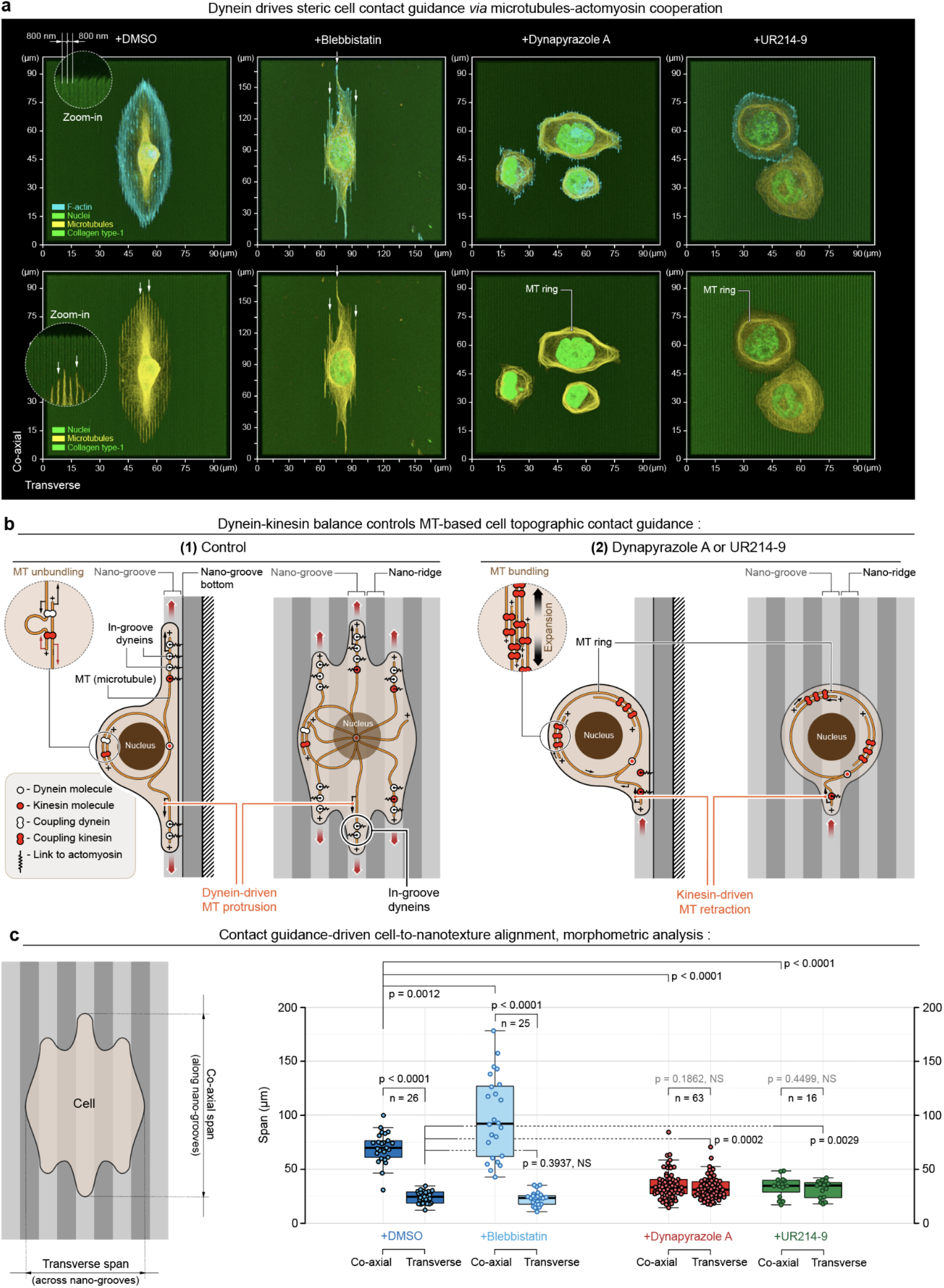
Dynein motor-driven in-groove MT guidance is crucial for the contact guidance-induced alignment of the cell to the “2.5D” collagen nanotexture. **(a)** - MDA-MB-231 cells (*top*) and their MT network (*bottom*) alignment to the sterically interactive collagen type-1-coated “2.5D” nanotextures (800/800 nm nano-grooves/nano-ridges, depth: 600 nm). *Left-*to*-right*: **(+DMSO)** - control conditions, **(+Blebbistatin)** - during actomyosin low contractility state, **(+Dynapyrazole)** - during dynein activity inhibition, and **(+UR214-9)** - during septin-9 actin-to-MT translocation and kinesin overactivation. Note that both direct dynein inactivation **(+Dynapyrazole A)** and kinesin outbalance over dynein activity **(+UR214-9)** suppress MT network-nanogrooves alignment, induces planar MT rings, and consequently suppresses cell contact guidance along the nanotextures. Note MTs-to-nanogroove alignment in active, balanced dynein cell states: **+DMSO** and **+Blebbistatin** (*arrows*); but MT network reorganize into rings with no alignment to the nanotexture during kinesins activity outbalance over dyneins: **+Dynapyrazole A** and **+UR214-9**. **(b)** - Schematic views of dynein+MT-driven cell contact guidance along the nanotextures **(1)**, and kinesin outbalance-driven loss of cell contact guidance **(2)** in either **+Dynapyrazole A** or **+UR214-9**. **(c)** - Quantification of cell contact guidance as cell spreading spans measurement along *vs*. across nanotextures shows no preferential response of cell spreading alignment to the nanotexture direction in both **+Dynapyrazole A** and **+UR214-9**. Pairwise one-way *t* tests-derived p values are shown on the plots with corresponding n (size of individual cells measurement sets) for each condition, generated in triplicates.

## CONCLUSIONS

Thus, mechanically and structurally antagonistic effects of dyneins and kinesins onto MT network as the cell scaffold are emerging as an additional and important cell mechanobiological system that requires a deeper insight. The inherent integration of the MT-actomyosin systems together *via* their motor proteins’ synergy appears as another dimension of the fundamental principles that regulate cell-microenvironment mechanical and structural integration and should find both theoretical and practical applications in cell biology, cell and tissue bioengineering, and medicine.

## ACKNOWLEDGEMENTS

E.D.T. and this work were supported by the Department of Pharmacology, Penn State College of Medicine *via* the startup funds. A.S.Z. and A.M. were supported by the FDA Intramural Research Program of the Center for Biologics Evaluation and Research. We thank Christian Combs and Daniela Malide for the Light Microscopy Core support at the National Heart, Lung, and Blood Institute, NIH. Work in the Burnette lab was supported by a MIRA to D.T.B. (R35 GM125028), a training grant to J.B.H. (T32 GM08320) and an American Heart Association fellowship to J.B.H. (AHA 836090). A.C.R. and C.S. were supported by the National Institutes of Health (NIH) Intramural Research Program in the National Institute of Biomedical Imaging and Bioengineering (NIH grant # ZIA EB000094) and by the NIH Distinguished Scholars Program.

## DATA AVAILABILITY STATEMENT

The authors declare that the data supporting the findings of this study are available within the paper, the Supporting Information, and in the Source Data file, and any data can be made further available upon reasonable requests.

## AUTHORS CONTRIBUTIONS

E.D.T. observed MT rings induction and conceived, formulated, and theoretically conceptualized the project, E.D.T., and A.S.Z. developed the theoretical framework, designed the experimental strategies, A.C.R. developed the theoretical and experimental methods for dynamic live cell micromechanical probing, E.D.T., A.S.Z., and A.C.R. carried out the core mechanobiology, cell biology, and the microscopy studies, E.D.T. microfabricated the artificial micropatterned ECM platforms for the MT rings study. A.M., C.S., and A.C.R. performed the AFM analysis, R.K.S. designed, synthesized, purified, chemically tested and verified septins inhibitor UR214-9 and outlined septin-kinesin signaling axis framework, E.D.T., A.S.Z., J.B.H., D.T.B., A.M., and R.K.S. performed the UR214-9 experiments and data analysis, J.B.H. and D.T.B. performed the live cell MT ring formation video-microscopy, E.D.T., A.S.Z., and A.C.R. interpreted the data, all co-authors contributed to the manuscript editing, E.D.T. wrote the manuscript, E.D.T. oversaw all aspects of this study.

## MATERIALS & METHODS

### Cell experiments

Human MDA-MB-231 (ATCC® HTB-26™) and monkey COS-7 (ATCC® CRL-1651™) fibroblast cells were maintained in DMEM with 4.5 g/L D-glucose, L-glutamine,110 mg/L sodium pyruvate (Corning Cellgro®, Cat#10013CV) and 10% heat-inactivated FBS (HyClone®, Cat#SH30071.03H) at 37°C in 5% CO2. Mouse Do-11-10 thyoma cells (Sigma, Cat#85082301) were maintained in ImmunoCult™-XF T Cell Expansion Medium (STEMCELL™ Technologies Inc., Cat#10981) with the addition of Human Recombinant Interleukin 2 (STEMCELL™ Technologies Inc., Cat#78036.3) at 37°C in 5% CO2. We treated cells in glass-bottom 35 mm Petri dishes (MatTek Corp., Cat#P35G-1.5-14-C) using (-)-Blebbistatin enantiomer (Sigma, Cat#203391), Dynapyrazole A (Sigma, Cat#SML2127), Kinesore (Sigma, Cat#SML2361), septin inhibitor UR214-9 (synthesized by Rakesh K. Singh), or dimethyl sulfoxide (Sigma, Cat#472301), as indicated in the main text.

### Synthesis of UR214-9

Equimolar mixture of 4-amino-2,6-dichloropyridine (Tokyo Chemical Industries, Cat#A2369) and 2-fluoro-3-(trifluoromethyl)phenyl isocyanate (Alrdich, Cat#472182-2G) in dimethylformamide (DMF) was stirred and heated at 65°C in a sealed glass tube overnight. DMF was removed using a Buchi rotary evaporator. The crude reaction mixture was dissolved in a mixture of dichloromethane(DCM)+MeOH and purified by preparative thin-layer chromatography using Hexane:EtOAc (50:50) as eluent. The pure product band was scrapped off the glass plate and UR214-9 was stripped from the silica gel using DCM+MeOH through a sintered funnel. The solvent was evaporated using a Buchi rotary evaporator to obtain UR214-9 as an off-white powder. UR214-9 was dried overnight in a desiccator and analyzed by proton and carbon NMR and mass spectrometry. MS-APCI:[368.3, singlet,100%; 370.2,doublet,60%]

### High precision micropatterning

A detailed, step-by-step instructive protocol for high-fidelity micropatterning is described elsewhere ^73^. Briefly, micro-contact printing of the high precision micron-scale features is a challenging task due to the van-der-waals and capillary effects between the micro-stamp and the printed surface that provoke a collapse of the soft conventional PDMS micro-stamps onto the printed intermediate (glass) surface. In order to prevent the collapse effects we substituted regular PDMS micro-stamps with composite stamps: soft PDMS blocks, veneered with a submillimeter-thick hard PDMS (hPDMS) for non-collapsing high-definition printing surfaces ^69,70^. For the hPDMS preparation protocol, please see “hPDMS formulation” section. In order to cast the micro-printing surface, we utilized the commercially fabricated and pre-passivated silicon molding matrix (Minnesota Nano Center, University of Minnesota), and coated them with ≤0.5mm hPDMS by gentle spreading with soft Parafilm-made spatula (Hach, USA), cured at 70°C for 30 minutes and subsequently cast with regular PDMS to the layer final thickness of 8 mm (rPDMS; 1:5 curing agent/base ratio, Sylgard-184, Dow Corning). Cured (at 70°C for ∼1 hour) composite micro-stamps were peeled, and cut into 5×5 mm or 1×1 cm pieces and used as the ready-to-use micro-stamps. With composite micro-stamps, we printed collagen micropatterns on PAA as follows: First, we microprinted α-collagen-1 rabbit pAb (AbCam, Cambridge, UK, Cat#ab34710; RRID:AB_731684), conjugated with biotin, ((+)-biotin *N*-hydroxysuccinimide ester, Sigma-Aldrich, Cat#H1759; as specified by the commercial protocol) and a fluorescent tag (Alexa Fluor® succinimidyl esters, Invitrogen™, Molecular Probes®, Cat#A20000, Cat#A20003; as per commercial protocol) on a clean, prebaked intermediate coverglass (FisherFinest™ Premium Cover Glass; #1, Cat#12-548-5 P; 450 °C, baked overnight). Micro-stamps were coated with printed antibody at a concentration of 0.2 mg/mL in phosphate-buffered saline (PBS) by incubation for 40 min at 37 °C in a humid chamber. Micro-stamps were then rinsed in deionized water and dried under a jet of filtered air, nitrogen or argon immediately before use. Second, glass-bottom 35 mm Petri dishes (MatTek Corp., Ashland, MA, Cat#P35G-1.0-20-C) were activated with 3-(trimethoxysilyl)propyl methacrylate (Sigma-Aldrich, Cat#6514) for covalent crosslinking with PAA gels. Third, 5 μL of PAA premixes with 5% streptavidin–acrylamide (ThermoFisher, Cat#S21379) of the defined rigidity ^48^ were promptly sandwiched between the activated dish and the micropatterned coverglass immediately after adding a curing catalyst (aminopropyltriethoxysilane (APS)). In order to achieve desired rigidity of elastic PAA, we modulated concentrations for both 40% acrylamide (40% AA) base (BioRad, Cat#161–0140) and its cross-linking molecular chain, 2% bis-AA (BioRad, Cat#161–0142) as described elsewhere ^48^. Additionally, streptavidin–acrylamide (Thermo Fisher, Cat#S21379) was added to the final concentration of 0.133 mg/mL to enable PAA gels cross-linking with biotinylated proteins of interest. Briefly, for preparation of 50 µL of 50 kPa PAA gel premix, the following components were mixed: 15 µL of 40% AA: 14.40 µL of 2% bis-AA; 3.33 µL of 2 mg/mL streptavidin-AA; 5 µL of 10×PBS; 11.17 µL of deionized milli-Q water; 0.1 µL of TEMED; 1 µL of 10% APS. The cured PAA sandwiches were then incubated at room temperature in deionized water for 1 h for hypotonic coverglass release from PAA gel. After coverglass removal, dish-bound gels retained fluorescent α-collagen-1 Ab micropatterns. Fourth, the resulting micropatterns or nanopatterns were incubated with 1 mg/mL rat monomeric collagen-1 (Corning, NY, Cat#354249) in cold PBS (4 °C, overnight), rinsed, and used for experiments.

### hPDMS formulation

For hard PDMS (hPDMS) we mixed 3.4 g of VDT-731 (Gelest, Inc., Cat#VDT-731), 18 μL of Pt catalyst (Platinum(0)-2,4,6,8-tetramethyl-2,4,6,8-tetravinylcyclotetrasiloxane complex solution) (Sigma-Aldrich, Cat#479543) and one drop of cross-linking modulator 2,4,6,8-Tetramethyl-2,4,6,8 -tetravinylcyclotetrasiloxane (Sigma-Aldrich, Cat#396281). Next, immediately before use, we added 1 g of HMS-301 (Gelest, Inc., Cat#HMS-301) and thoroughly mixed it for 30 sec on a vortex mixer ^69,71^.

### PAA elastic gels premixes

We chose to control PAA mechanical rigidity *via* modulation of concentration for both 40% acrylamide (40% AA) base (BioRad) and its cross-linking molecular chain, 2% bis-AA (BioRad) as described elsewhere ^72,73^. Additionally, streptavidin-acrylamide (Thermo Fisher) was added to the final concentration of 0.133 mg/mL to enable PAA gels cross-linking with biotinylated proteins of interest. Briefly, for preparation of 50 µL of G’ = 2.3 and 50 kPa PAA gel premixes, respectively, the following components were mixed: 40% AA: 9.33 and 15 µL; 2% bis-AA: 1.88 and 14.40 µL; 2 mg/mL streptavidin-AA: 3.33 and 3.33 µL; 10X PBS: 5 and 5 µL; deionized milli-Q water: 30 and 11.17 µL; TEMED: 0.1 and 0.1 µL; 10% APS: 1 and 1 µL. The premix solutions were degassed and stored at 4°C before use.

### Super- and high-resolution and confocal imaging

We fixed cells with cold (+4°C) DMEM with 4% Paraformaldehyde (P6148, Sigma) for 15 minutes, followed by two cycles of rinsing in 1% BSA in PBS, and 30 minutes of blocking-permeabilization using 0.1% Triton X-100 (X100, Sigma) - 10% BSA (BP9704, Fisher) in PBS (10010023, Thermo Fisher). For staining, we used β-tubulin (Sigma, Cat#T7816), α-tubulin (Abcam, Cat#ab18251), dynein (EMD Millipore, Cat#MAB1618), kinesin (EMD Millipore, Cat#MAB1613), septin-9 (Sigma, Cat#HPA042564) and septin-2 (Clonentech, Cat#60075) antibodies in 1% BSA PBS for 1 hour at room temperature. All labelings using Alexa-Fluor secondary antibodies (Thermo Fisher) were performed at their final concentration of 5 µg/mL for 1 hour in 1% BSA PBS at room temperature. After washing out the excess secondary antibodies, F-actin was stained with fluorescent phalloidin (Alexa Fluor phalloidin conjugates, Thermo Fisher Scientific; 10 U/mL). Chromatin was labeled with 1:1000 Hoechst solution (Tocris, Cat#5117). We mounted samples using 90% Glycerol (Sigma, Cat#G5516) in PBS.

High-resolution 3D and 2D imaging for cell morphometric analysis were performed on a Leica TCS SP8 laser scanning confocal microscope with LIAchroic Lightning system and LAS X Lightning Expert super-resolution capacity, 405, 488, 552 and 638 nm excitation diode lasers, with 40×/1.3 oil immersion objective (Leica, Germany). The scanning settings were optimized with Nyquist LAS X function with HyD2.0-SMD excitation sensors, at the regular pinhole size of 0.85 AU, and the scanning frequency at 100 Hz.

Alternatively, Nikon TiE stand with an A1Rsi Confocal scan head, powered by NIS-Elements Confocal software (Nikon, Japan) used for cell imaging. Objectives used were PlanApo VC 20×/0.75 NA and PlanApo VC 60×WI/1.20NA and excitation was provided sequentially by 405 nm, 488 nm and 561 nm lasers. Fluorescence was collected through a 1.2 AU pinhole using emission filters of 425-475nm, 500-550nm, and 570-620nm. Pixel size was adjusted to Nyquist sampling (voxel size x,y,z for the 20× objective, j,k,l for the 60x objective).

Morphometric analysis was performed automatically and/or manually utilizing LAS X (Leica, Germany) or NIS-Elements Advanced Research software (Nikon, Japan) as an integral part of the data analysis streamline “microscopy-to-measurement-to-analysis”. Video-sequences were also analyzed with ImageJ/FIJI “stacks” plug-in. Additionally, live cell imaging microscopy experiments were performed on an Olympus X81 (Olympus, Japan) microscopy inside a temperature (37°C), CO_2_ (5%), and humidity controlled chamber at 20× magnification. Brightfield and fluorescent images were obtained every 2 min using MetaMorph software (Molecular Devices, USA). Composite 2D/3D cells+micropattern images were reconstructed and assembled using NIS-Elements AR and linear image parametric adjustments. Figures were composed using unmodified NIS-Elements AR-generated TIFF images with Adobe Illustrator CC 2017 (Adobe).

Instant structured illumination microscopy (iSIM) was performed using the instant structured illumination microscopy (iSIM) by an Olympus IX-81 microscope (Olympus, Corp., Tokyo, Japan) equipped with an Olympus UPLAPO-HR ×100/1.5 NA objective, two Flash-4 scientific CMOS cameras (Hamamatsu, Corp., Tokyo, Japan), an iSIM scan head (VisiTech Intl, Sunderland, UK), and a Nano-Drive piezo Z stage (Mad City Labs, Madison, WI). The iSIM scan head included the VT-Ingwaz optical de-striping unit (VisiTech Intl, Sunderland, UK). Image acquisition and system control was done using MetaMorph Premiere software (Molecular Devices, LLC, San Jose, CA). Images were deconvolved with an iSIM-specific commercial plugin from Microvolution (Cupertino, CA) in FIJI.

### Live cell imaging of giant microtubule rings

Human MDA-MB-231 cells were transferred from confluent T25 flasks (CytoOne; Cat# CC7682-4325) into a 24-well plate (CytoOne; Cat# CC7682-7524) at 40% confluency. Cells were incubated overnight with 100nM SiR-Tubulin (Cytoskeleton, Inc.; Cat.#CY-SC002), then seeded at high density onto micropatterned coverslips (same as for immunofluorescence; see above) in DMEM containing 100nM SiR-Tubulin. After either 30 minutes or 1 hour of spreading, media was replaced with DMEM containing 100nM SiR-Tubulin, 130uM Dynapyrazole A, and 100uM Blebbistatin. Cells were imaged using a Visitech “instant” Structure Illumination Microscopy (iSIM) microscope and images were captured either every 10 minutes or every 30 minutes in the far-red channel using a CMOS camera.

### Preparation of collagen-1-functionalized acrylamide gels

We used composite stamps made of hard polydimethylsiloxane (hPDMS) base and regular PDMS support. For hPDMS, we mixed 3.4 g of VDT-731 (Gelest, Inc.), 1 g of HMS-301 (Gelest, Inc.), 18 µL of Pt catalyst (Platinum(0)-2,4,6,8-tetramethyl-2,4,6,8-tetravinylcyclotetrasiloxane complex solution, Sigma) and one drop of cross-linking modulator (2,4,6,8-Tetramethyl-2,4,6,8-tetravinylcyclotetrasiloxane, Sigma). We thoroughly mixed hPDMS for 30 seconds on a vortex mixer, degassed it using centrifugation at 400 g for 3 minutes, immediately used it for SU8 mold coating, and cured at 65°C for 1 hour. For the supporting layer, we prepared regular PDMS as 1:10 curing agent/base ratio mixture formulation (Sylgard-184, Dow Corning), removed air bubbles by centrifugation at 400 g for 3 minutes, coated hPDMS, and cured at 65°C for 4 hours. We cut peeled PDMS replicas into individual stamps using a sharp blade.

We did antibody labelling following commercial protocols for (+)-Biotin N-hydroxysuccinimide ester (H1759, Sigma) and Alexa Fluor™ 488 NHS Ester (A20000, Thermo Fisher) products. We coated individual PDMS stamps with biotin-Alexa labelled Anti-Collagen I antibody (Abcam, Cat#ab34710) in PBS solution (0.1 mg/mL, in a wet chamber, overnight at 4°C). After the incubation with the antibody solution, we gently rinsed stamps in deionized water, and blow dried them under a filtered air.

For making flat collagen grids, we first printed anti-collagen-I antibody patterns onto the “intermediate” 2D glass surfaces. For making grooves, we used antibody-coated 2.5D stamps directly. Biotinylated antibodies on intermediate glass surfaces or stamps were cross-linked to polymerizing PAA gels. For that we polymerized 7-10 μL of gel mix in the “sandwich” fashion between a patterned surface and glass-bottom 35 mm Petri dishes (MatTek Corp., Ashland, MA), that we activated with 3-(trimethoxysilyl) propyl methacrylate (Sigma-Aldrich) in ethyl alcohol (Pharmco-Aaper) and acetic acid (Fisher Chemical) as per the commercial protocol. For preparation of G’= 50kPa gels, we mix degassed 40% acrylamide: 15 µL and 2% bis-acrylamide: 14.4 µL; with 2 mg/mL streptavidin-acrylamide: 3.33 µL; 10X PBS: 5 µL; deionized milli-Q water: 11.17 µL; TEMED: 0.1 µL; 10% APS: 1 µL.

### Atomic Force Microscopy

Suspended MDA-MB-231 triple negative breast cancer cells with passage number between P8-P13 were plated on a glass-bottom dish (Willco Wells) precoated with 0.01% poly-L-lysine (Sigma-Aldrich) with culture media solution (Life Technologies). Depending on the pharmacological drug treatment, cells were incubated with times between ∼15 min to 75 min to let them weakly adhere to the glass-bottom dish surface and the drug to take maximum effect. AFM force spectroscopy experiments were performed using a Bruker BioScope Catalyst AFM system (Bruker) mounted on an inverted Axiovert 200M microscope (Zeiss) equipped with a confocal laser scanning microscope 510 Meta (Zeiss) and a 40x objective lens (0.95 NA, Plan-Apochromat, Zeiss). The combined microscope instrument was placed on an acoustic isolation table (Kinetic Systems). During AFM experiments, cells were maintained at physiologically relevant temperature 37°C using a heated stage (Bruker). A soft silicon nitride tipless AFM probe (HQ:CSC38/tipless/Cr-Au, MikroMasch) was used for gentle compression of spherical weakly-adhered 231 cells. The AFM microcantilevers were pre-calibrated using the standard thermal noise fluctuations method, with estimated spring constants for microcantilevers used between 0.05–0.11 N/m. Immediately after probe pre-calibration, the AFM tipless probe was moved on top of a spherical MDA-MB-231 cell. Five to ten successive force curves were performed on each cell. The deflection set-point was set between 12-32 nm yielding applied forces between 0.95 to 2.7 nN.

All AFM force-distance curves measurements were analyzed using a custom-written MATLAB (The MathWorks) code to calculate the cellular surface cortical tension. For curve fitting, indentation depths between 0-600 nm were relatively consistent in yielding good fits (R2>0.75). Curves with poor fits R2<0.75 were discarded from the analysis. Additionally, we discarded noisy force curves and/or curves that presented jumps possibly due to cantilever and plasma membrane adhesion, slippage, or very weakly adherent moving cells.

Spherical 231 cells cellular surface cortical tension (T; pN/µm) was calculated by fitting each recorded force-distance curve using the cortical tension method we previously described that defines the force balance relating the applied cantilever force with the pressure excess inside the rounded cells and the corresponding actin cortex tension; 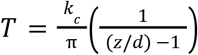
, where T is the cellular surface tension, kc is the AFM cantilever spring constant, Z is the Z-piezo extension, and d is the cantilever mean deflection ^74^.

## Statistical analysis

Only pairwise comparisons as one-sided *t* tests between a control group and all other conditions are utilized to analyze the data, as well as between paired -Blebbistatin and +Blebbistatin co-treatment groups. Statistical analysis is performed using either KaleidaGraph 4.5.4 (Synergy Software) or Prism 7b (GraphPad Software, Inc). The exact p values are indicated on the plots, unless the p<0.0001, *i*.*e*., below cut-off lower limit for Kaleidagraph and Prism softwares. Sample size n for each comparison is reported in the corresponding plots (*i*.*e*., n reflects the number of measured individual cells). Number of replicates is 3, unless specified otherwise. Data are shown as box and whiskers diagrams: first quartile, median, third quartile, and 95% percent confidence interval.

## Supporting Information

**SI1.**
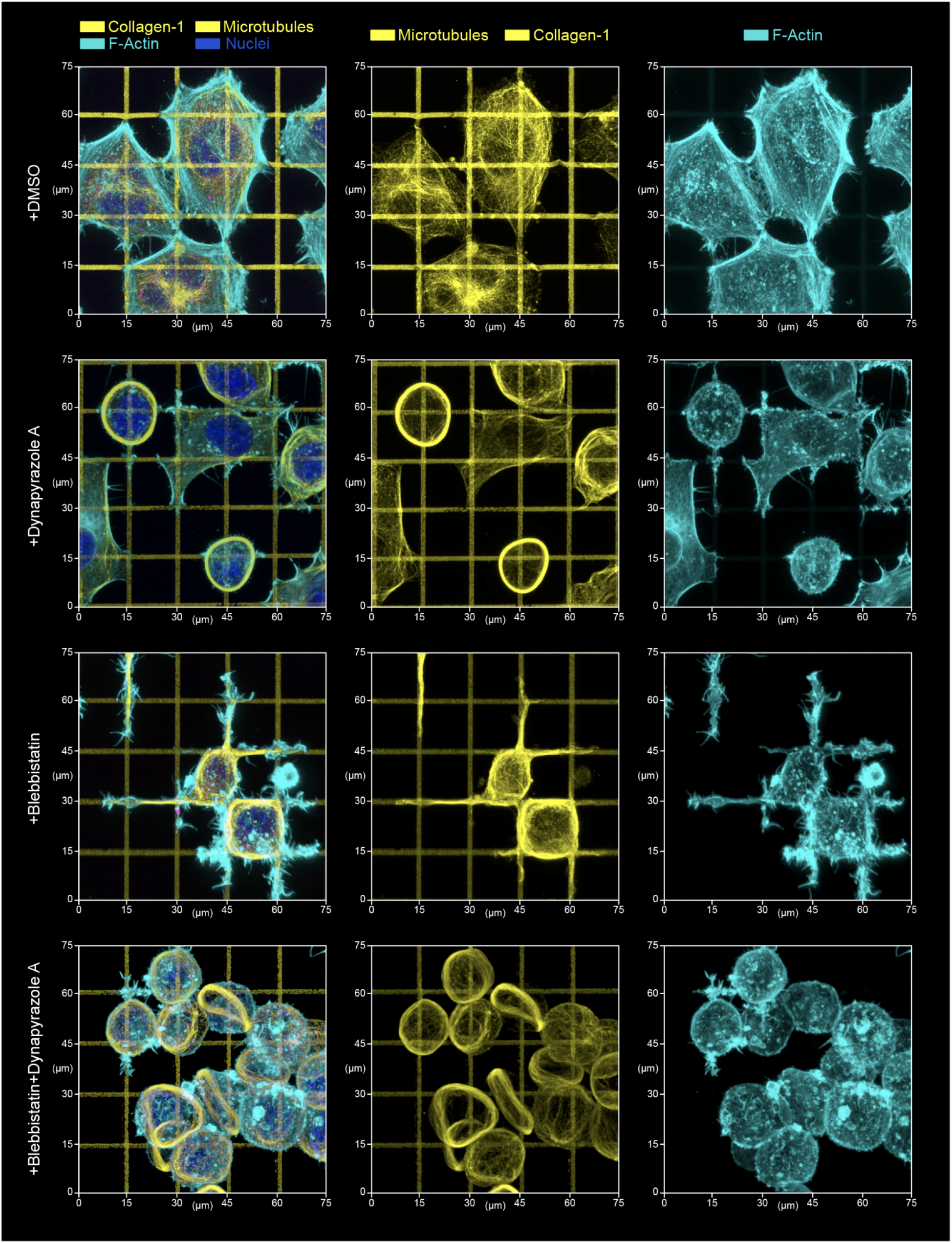
Top 3D overview of MDA-MB-231 cells during their adhesion and spreading along the artificial extracellular matrix (orthogonal collagen grid, G’=50 kPa). Corresponds to Figure 1a. Micrographs present cell configurations in the control condition **(+DMSO)**, during dynein inhibition **(+Dynapyrazole A)**, in the presence of actomyosin contractility inhibitor **(+Blebbistatin)**, and under combined dynein and actomyosin contractility suppression **(+Blebbistatin+Dynapyrazole A)**. Upon biaxial spreading along the orthogonal collagen lanes in the control conditions **(+DMSO)** cells develop a polygonal architecture with stress-fibers predominantly developing at the mechanically tensed cell periphery - *i*.*e*., concave free cell edges. Suppression of the dynein motor protein activity **(+Dynapyrazole A)** results in cell spreading architecture disarray, reduced polygonal cell architectures and partially ‘dendritic’-like cell morphologies (see also **Figure 3c, +Dynapyrazole A**). Actomyosin contractility inhibition **(+Blebbistatin)** shifts cell cytoskeletal organization towards the spreading network of dendritic protrusions that compliantly extend along the collagen grid with high cell dendrites-to-grid structural fidelity. These dendritic protrusions have the microtubule scaffold as their core. Notably, combined dynein and myosin II inhibition **(+Blebbistatin+Dynapyrazole A)** results in the suppressed cell spreading (but not adhesion), and formation of MT rings that deform non-spread spheroid cells into the lenticular discoids.

**SI2.**
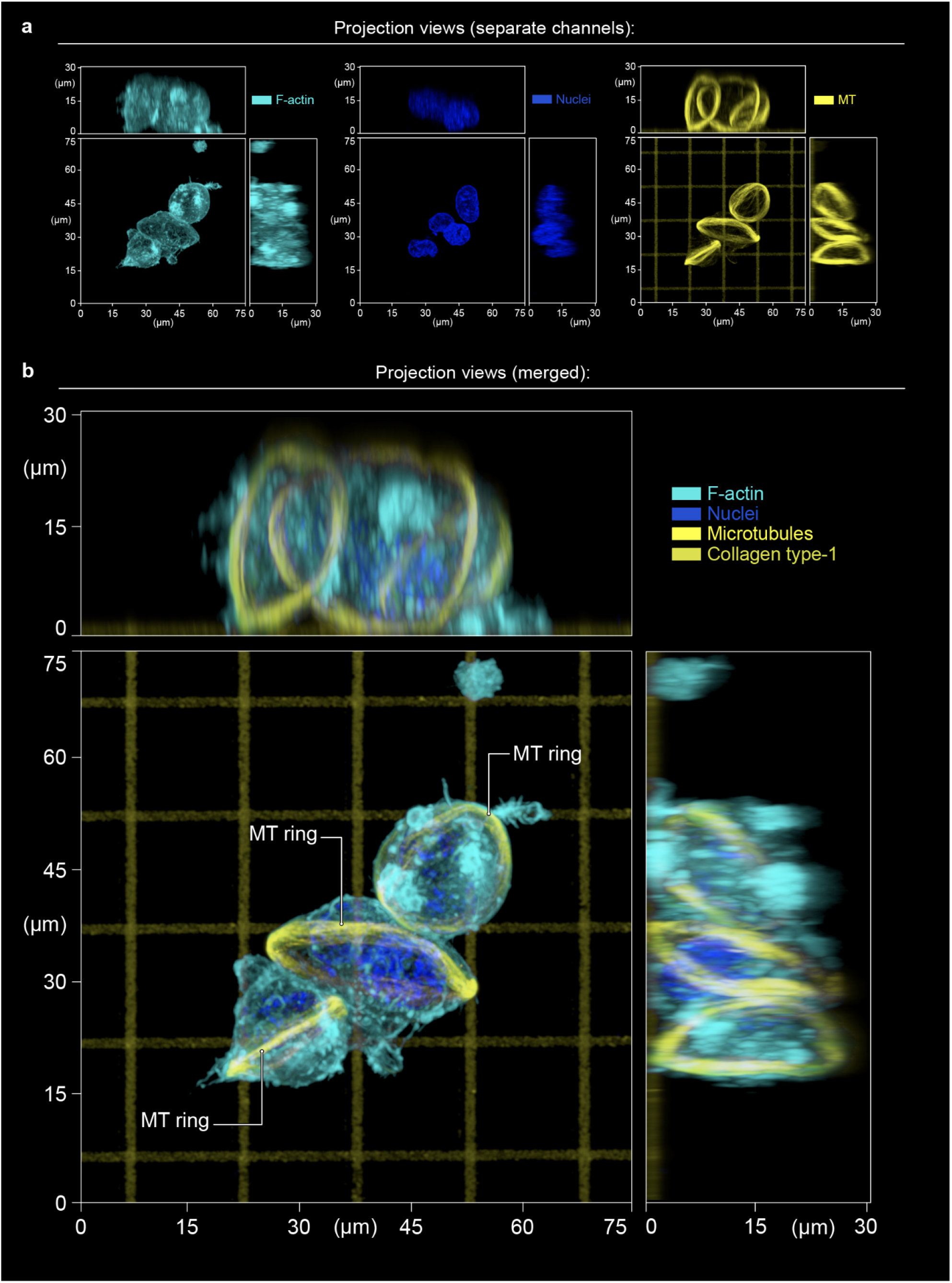
Detailed projection views of discoid MDA-MB-231 cells that were formed during +Dynapyrazole A+Blebbistatin treatment (corresponds to Figure 1b and 1c). **(a)** - Separate channel 3D projection views: F-actin (phalloidin), nuclei (Hoechst), microtubules (tubulin immunofluorescence imaging), collagen type-1 micropatterned grids (prelabeled). **(b)** - Enlarged and detailed 2D projection views of the merged channels.

**SI3.**
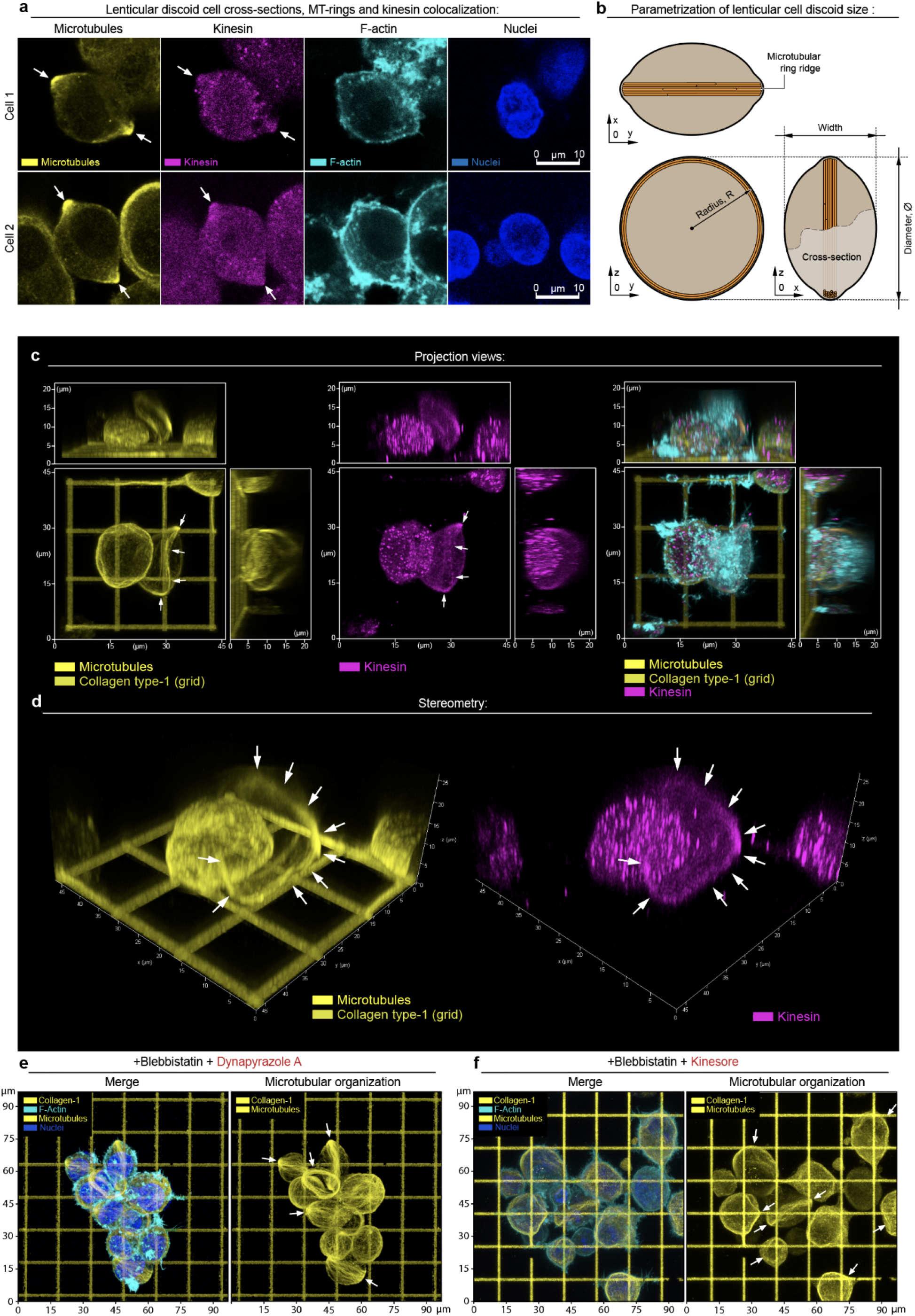
Details of structural dissection of the MT ring composition. **(a)** - Detailed channel-separated views of discoid cell median cross-sections (*cell 1 and cell 2*). Corresponds to **Figure 3a**. **(b)** - Detailed schematic of the discoid cell structural parameters measurement (width *vs*. diameter). Corresponds to **Figure 3b**. **(c, d)** - Standard 2D projection views **(c)**, and stereometric views of the MT ring and the kinesin immunofluorescent spatial distribution **(d)**, showing kinesin colocalization with the **+Dynapyrazole A+Blebbistatin**-induced MT rings (*arrows*). Corresponds to **Figure 3a**. **(e, f)** - Detailed, channel-separated view of the MT rings (*arrows*) in **+Dynapyrazole A+Blebbistatin (e)**, as compared to the alternative kinesin outbalance induction *via* **+Kinesore+Blebbistatin** treatment **(f)**. MT rings are outlined by arrows. Corresponds to **Figure 3e**.

**SI4.**
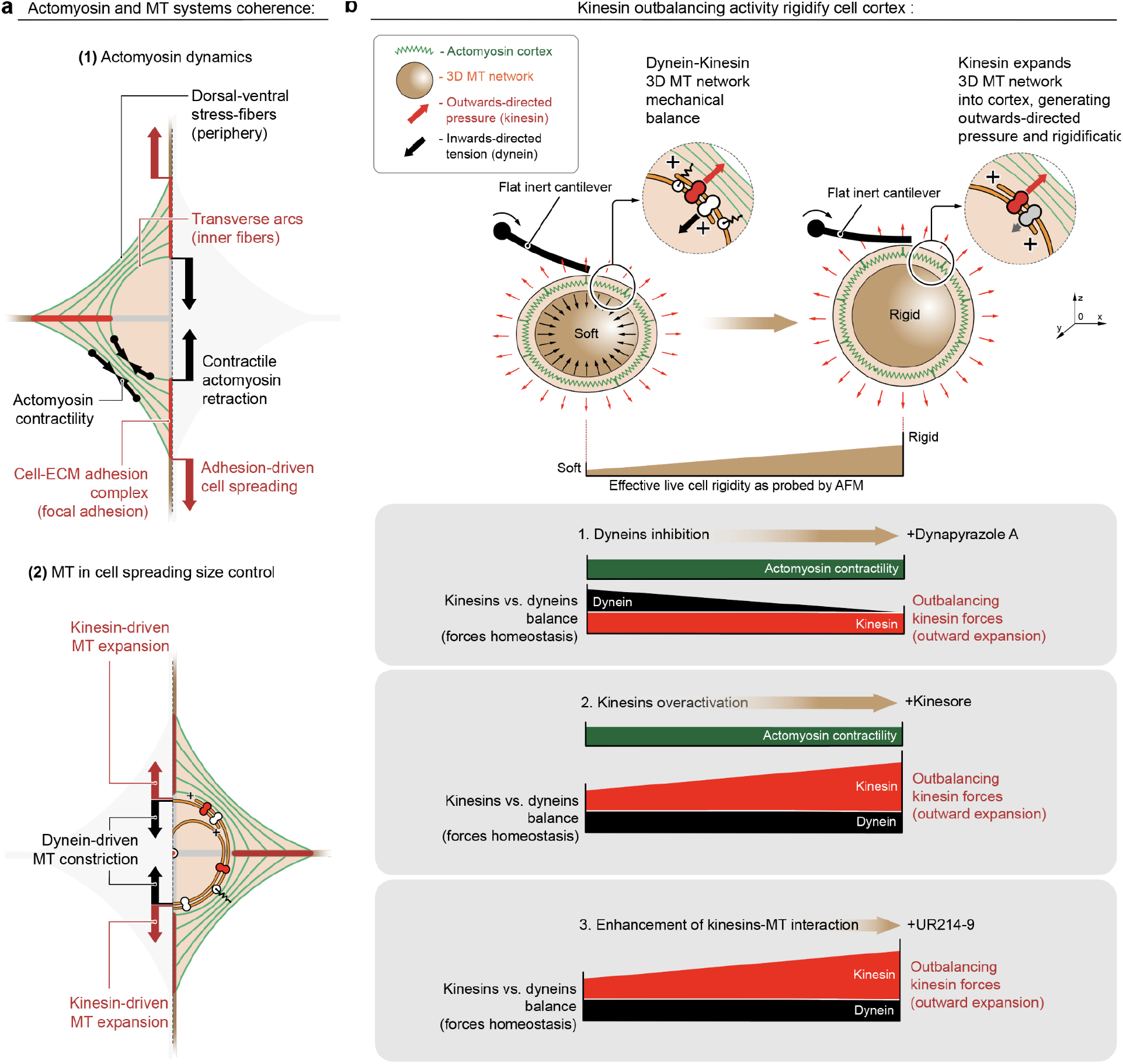
Dynein-kinesin balance controls cell size during spreading, as well as cell rigidity. **(a)** - MDA-MB-231 cell spreading on collagen-1 orthogonal grid (biomimetic ECM) is a result of mechanical superposition of **(1)** contractile actomyosin cytoskeleton adhesion system (stress-fibers+focal adhesions), and **(2)** collagen “fibers” organize the cell mechanical dynamics into a balanced cell spreading process and cell contractile MT network expansion-contraction, driven by kinesin-dynein counterbalancing activity. Corresponds to **Figure 3d**. **(b)** - Alternative cell rigidification mechanisms: (*Top*) - Schematics of the cell rigidification, driven by kinesin-wise activity outbalance over dyneins that leads to the kinesin-driven mechanical expansion of the MT network into the cell cortex. (*Bottom*) - **(1)** Dynein inhibition with Dynapyrazole A, Kinesin overactivation with Kinesore **(2)**, as well as with septin-9-MT complex, induced by UR214-9 septin filaments disassembly **(3)** all lead to the dynein-kinesin activity balance shift towards kinesins, leading to the similar final effects of the cell cortex rigidification, detected by AFM. Corresponds to **Figure 4c**.

**SI5.**
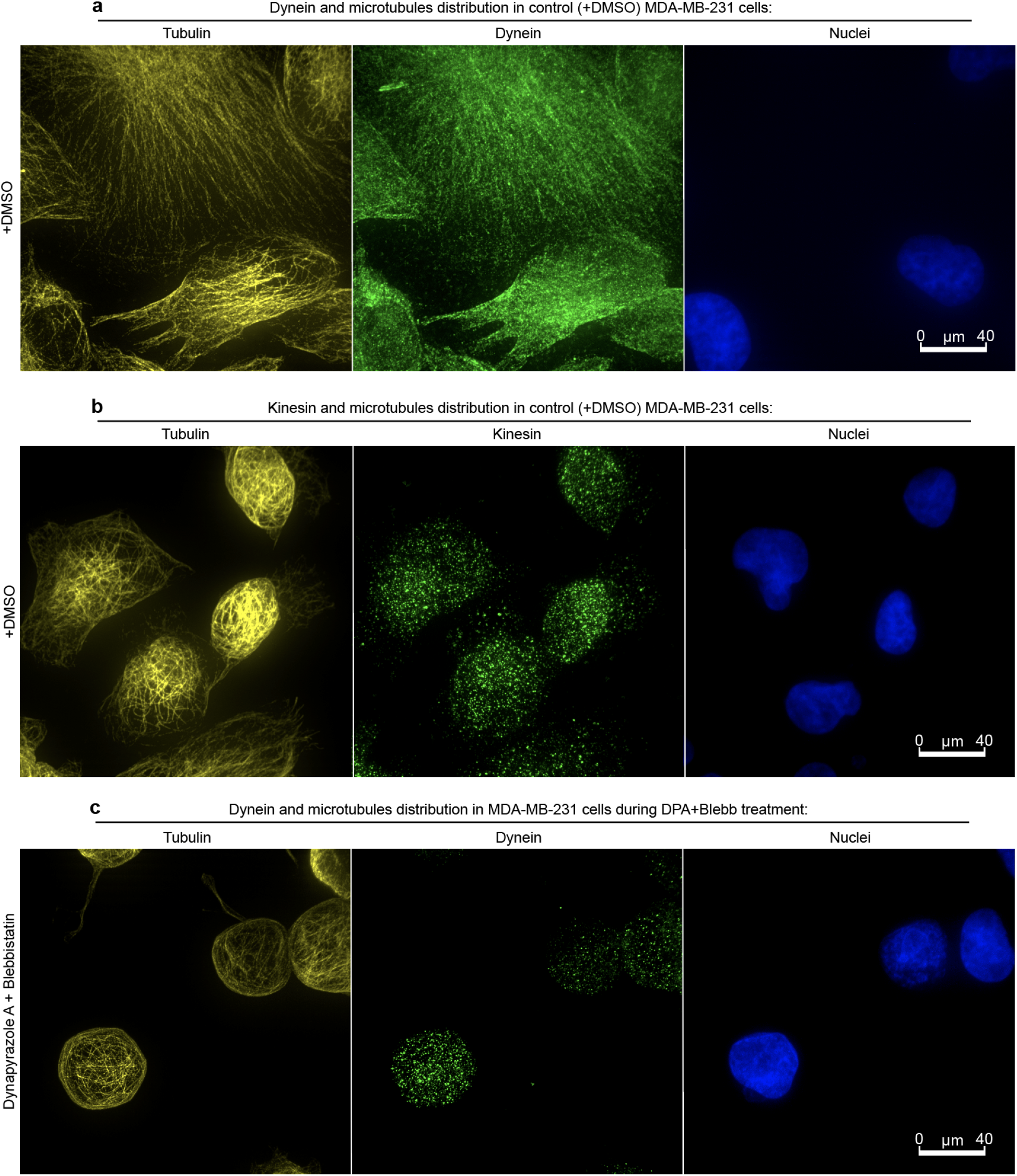
Localization of dyneins and kinesins in control conditions (+DMSO), and during +Dynapyrazole A+Blebbistatin treatments. **(a)** - Dyneins colocalize with the MT network in **+DMSO**. Corresponds to **Figures 3a, 4b**. **(b)** - Kinesins show diffuse distribution throughout cell volume in DMSO, but shows strong MT ring colocalization in **+Dynapyrazole A+Blebbistatin**, see **Figure 3a** and **SI3a, SI3c, and SI3d**. Corresponds to **Figure 3a, 4b**. **(c)** - **Dynapyrazole A+Blebbistatin** treatment induces dynein-MT dissociation, dynein translocation from MT network into cytoplasm. Corresponds to **Figure 3a**.

**SI6.**
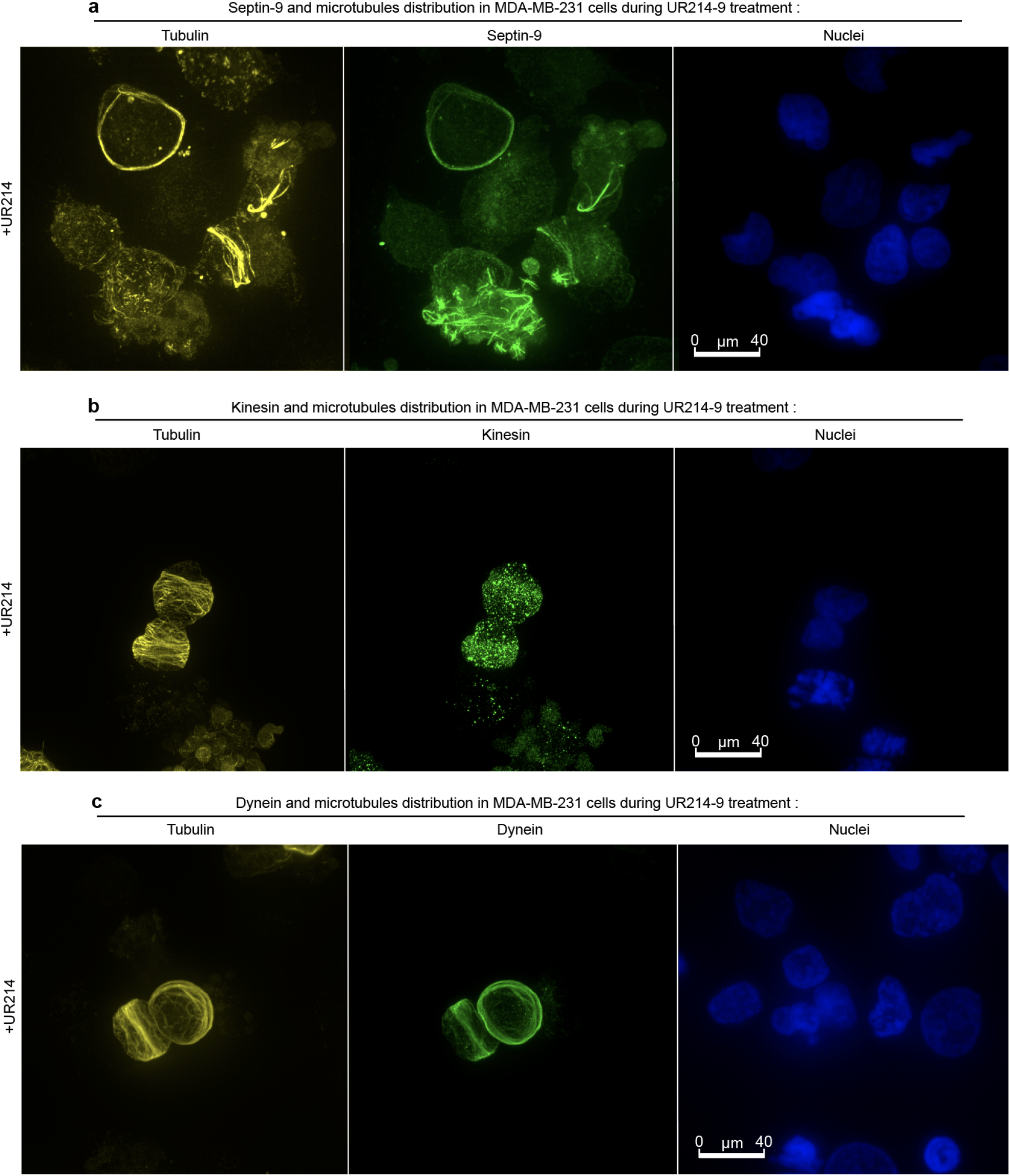
Dyneins, kinesins and septin-9 localization in MDA-MB-231 cells during UR214-9 treatment. **(a)** - Septin-9 localizes at MT network in **+UR214-9** conditions. **(b)** - Kinesins colocalize with forming MT rings during microtubule translocation of septin-9 in UR214-9 treatment. **(c)** - Dyneins colocalize with MT network during UR214-9 treatment, since UR214-9 overactivates kinesins, but does not suppress dyneins. Corresponds to **Figures 4b, 4d**, and **4e**.

